# Motility-dependent processes in *Toxoplasma gondii* tachyzoites and bradyzoites: same same but different

**DOI:** 10.1101/2024.09.28.615543

**Authors:** Robyn S. Kent, Gary E. Ward

## Abstract

The tachyzoite stage of the apicomplexan parasite *Toxoplasma gondii* utilizes motility for multiple purposes during its lytic cycle, including host cell invasion, egress from infected cells, and migration to new uninfected host cells to repeat the process. Bradyzoite stage parasites, which establish a new infection in a naïve host, must also use motility to escape from the cysts that are ingested by the new host and then migrate to the gut wall, where they either invade cells of the intestinal epithelium or squeeze between these cells to infect the underlying connective tissue. We know very little about the motility of bradyzoites, which we analyze in detail here and compare to the well-characterized motility and motility-dependent processes of tachyzoites. Unexpectedly, bradyzoites were found to be as motile as tachyzoites in a 3D model extracellular matrix, and they showed increased invasion into and transmigration across certain cell types, consistent with their need to establish the infection in the gut. The motility of the two stages was inhibited to the same extent by cytochalasin D and KNX-002, compounds known to target the parasite’s actomyosin-based motor. In contrast, other compounds that impact tachyzoite motility (tachyplegin and enhancer 5) have less of an effect on bradyzoites, and rapid bradyzoite egress from infected cells is not triggered by treatment with calcium ionophores, as it is with tachyzoites. The similarities and differences between these two life cycle stages highlight the need to characterize both tachyzoites and bradyzoites for a more complete understanding of the role of motility in the parasite life cycle and the effect that potential therapeutics targeting parasite motility will have on disease establishment and progression.

## Introduction

*Toxoplasma gondii* is a prevalent human pathogen, infecting >30% of the global human population ^[1-4]^. Infection with *T. gondii* mostly occurs via ingestion of either bradyzoite-containing tissue cysts in undercooked meat, or sporozoite-containing oocysts in cat feces that can also contaminate water sources ^[5]^. While acute infection is often subclinical in immune competent individuals, ocular ^[6, 7]^ and pulmonary ^[8]^ disease can occur, particularly associated with virulent atypical strains ^[6]^.

Bradyzoites, after excysting from tissue cysts, must efficiently migrate to the wall of the gut epithelium and either actively invade the cells of the gut or penetrate between these cells to access the lamina propria ^[9]^. Following infection, the bradyzoites transition into the fast-replicating tachyzoite stage. The tachyzoites disseminate throughout the body and, following repeated rounds of invasion, replication and egress from host cells, they elicit and are largely cleared by the host immune response ^[10]^. However, a subset of the tachyzoites differentiate into bradyzoites and build a protective cyst wall, primarily in the brain and skeletal muscle, to establish a chronic infection ^[11]^. Bradyzoites replicate much more slowly than tachyzoites and have traditionally been considered more dormant ^[12, 13]^ although recent data show the cysts grow and parasites within replicate *in vivo* ^[14]^. The cysts are resistant to all current therapeutics ^[15]^ and can establish an infection in a new host if ingested ^[11]^. Cysts can also reactivate *in situ* if the host becomes immune compromised ^[16, 17]^, causing potentially life-threatening disease.

Much is known about motility of the tachyzoite stage ^[18-20]^, including the composition of the motor complex thought to power motility and the role of several proteins that link the parasite motor to the substrate along which the parasite moves ^[21-25]^. Furthermore, we know that tachyzoite motility is not only important for parasite movement between cells and through tissues, but also for other essential aspects of the lytic cycle such as invasion into and egress from host cells ^[24, 26]^. In contrast, very little is known about motility and motility-dependent processes in the bradyzoite stage, although motility is clearly critical for establishing infection after the ingestion of tissue cysts. Tachyzoites are often referred to as the “highly motile” stage, a viewpoint consistent with the recent report that bradyzoites chemically liberated from cysts show reduced motility in 2D assays compared to mechanically liberated tachyzoites ^[27]^. A more systematic comparison of the motile behavior and motility mechanisms of these two life-cycle stages will shed new light on the biological processes underlying *Toxoplasma* infection.

We show here that, after excystation using a brief protease treatment and mechanical shear, bradyzoites are in fact as motile within a model extracellular matrix as tachyzoites, show enhanced invasion into two gut cell lines and surpass tachyzoites in their ability to transmigrate across certain polarized epithelial cell monolayers. Bradyzoites respond similarly to tachyzoites when treated with some modulators of tachyzoite motility, and differently to others. Finally, we confirm previous work showing that – in contrast to tachyzoites – calcium stimulation does not lead to simultaneous, rapid egress of bradyzoites from infected host cells. Together, these data demonstrate that while there are similarities in the motility of tachyzoites and bradyzoites, there are also stage-specific differences that will need to be considered in current efforts to therapeutically target motility-dependent processes ^[28-32]^ in these widespread human and animal pathogens.

## Methods

### Cell and parasite culture

Tachyzoite stage parasites (RH, PRU-RFP and PRU-LDH2GFP) were propagated by serial passage in a primary fibroblast-like cell line from human foreskin provided by Dr. Thomas Moehring. HFFs were grown to confluence in Dulbecco’s Modified Eagle’s Medium (DMEM) (Life Technologies, Carlsbad, CA) containing 10% v/v heat-inactivated fetal bovine serum (FBS) (Life Technologies, Carlsbad, CA), 10 mM HEPES pH 7, and 100 units/ml penicillin and 100 μg/ml streptomycin, as previously described ^[33]^. Prior to infection with *T. gondii*, the medium was changed to 1% DMEM, DMEM supplemented with 10mM HEPES pH 7, 100 units/ml penicillin and 100 μg/ml streptomycin, and 1% v/v FBS. Differentiation to bradyzoite stage parasites was induced by replacing the media 1 – 2 hours after challenge with tachyzoites to differentiation medium (RPMI supplemented with 10mM HEPES pH 8.2, 100 units/ml penicillin and 100 μg/ml streptomycin, 2mM l-glutamine and 1% v/v FBS) and grown in ambient CO2 conditions, as previously described ^[34]^. For all *in vitro* bradyzoite assays the differentiation medium was replaced daily for a minimum of 5 days before parasite isolation. The cell lines used for invasion and transmigration assays were authenticated as follows: authenticated Hs27 human foreskin fibroblasts (HFFs), HCT-8 cells and EA.Hy926 cells were purchased directly from American Type Culture Collection (ATCC); MDCK cells were authenticated by short tandem repeat (STR) profiling by IDEXX (Westbrook, ME); and CACO-2 cells were authenticated by STR profiling at the Vermont Integrative Genomics Resource (Burlington, VT).

### Mouse infections

Infections were carried out in 4–5-week-old male CBA/J mice purchased from Jackson Laboratories (Bar Harbor, ME). All mice were socially housed and acclimated for at least 3 days prior to infections. Mice (two mice per biological replicate were pooled per motility experiment) were challenged by intraperitoneal injection of 100 PRU-LDH2GFP parasites resuspended in a total volume of 200μl sterile PBS. Daily animal health monitoring began two days after infection. Mice were euthanized if predetermined welfare thresholds were exceeded.

### Isolation of bradyzoites from infected mice

Parasites were harvested for assays 28 – 32 days after infection. Brains were washed in 5ml PBS before homogenization in PBS (2ml/brain) using a Dounce homogenizer. Two brains per sample were pooled before serially passing through 18G, 20G and 22G needles five times, and centrifuged at 2000*xg* for 10 minutes. Each sample was thoroughly resuspended in 6ml 20% dextran-150 in PBS and centrifuged at 400*xg* for 10 minutes. The myelin layer and dextran-150 were removed by aspiration. The pellet was washed once with 10ml PBS by centrifugation at 2000*xg* for 10 minutes. To excyst the bradyzoites, the pellet was resuspended in 1ml acid pepsin (AP) stock solution (0.01 mg/mL in 1% NaCl pH 2.1) diluted in 1ml freshly made 1% NaCl and incubated at 37°C for 1 minute. The suspension was then passed through a 26G needle, directly into 2ml Na_2_CO_3_ and 2ml 1% DMEM to inactivate the pepsin. The released parasites were filtered through a 3-μm Whatman Nuclepore filter (Millipore Sigma), pelleted (5 min at 2000*xg*) and resuspended in motility buffer (1× Minimum Essential Medium lacking sodium bicarbonate, 1% (v/v) FBS, 10 mM HEPES pH 7.0 and 10 mM GlutaMAX L-alanyl-L-glutamine dipeptide as in ^[20]^(for motility assays) or 1% DMEM (for invasion and transmigration assays) as outlined below.

### Isolation of bradyzoites from cell culture

Bradyzoites were isolated by passing infected cells through a 22G needle, centrifugation (2000*xg*, 5 minutes), resuspension in 1ml pepsin (0.01 mg/mL in 1% NaCl pH 2.1) and 1ml 1% NaCl, and incubation at 37°C for 1 minute. Mechanical disruption of the cyst wall was facilitated by passage through a 26G needle. The pepsin was inactivated with 2ml Na2CO3 and 2ml 1% DMEM and the parasites were then filtered through a 3-μm Whatman Nucleopore filter (Millipore Sigma). Bradyzoites were pelleted (5 min at 2000*xg*) and resuspended in motility buffer ^[20]^ (for motility assays) or 1% DMEM (for invasion and transmigration assays) as outlined below.

### Isolation of tachyzoites from cell culture

Tachyzoites were isolated from infected HFF cells by syringe release through a 26G needle, filtering through a 3-μm Whatman Nucleopore filter (Millipore Sigma), centrifugation (2000*xg*, 5 minutes) and resuspension in motility buffer (for motility assays) ^[20]^ or 1% DMEM (for invasion and transmigration assays) as outlined below.

### Parasite motility assays

3D motility assays were performed as previously described ^[20]^. 1024 pixel × 384 pixel images were captured over 80 sec using an iXON Life 888 EMCCD camera (Andor Technology) driven by NIS Elements software v.5.11 (Nikon Instruments). Image stacks consisted of 41 z-slices, captured 1 um apart with a 16ms exposure time. Tachyzoites were tracked using Hoechst 33242 fluorescence and bradyzoites were tracked using cytosolic GFP fluorescence. Tracking was achieved with Imaris ×64 v. 9.2.0 (Bitplane AG, Zurich, Switzerland). The minimum displacement cutoff thresholds were determined using heat-killed parasites (as previously described ^[20]^ to be 2.5μm and 2.8μm for tachyzoites and bradyzoites, respectively.

### Invasion Assays

The invasion of tachyzoites (PRU-RFP) and bradyzoites (PRU-LDH2GFP) was measured in co-invasion assays, where both stages were applied to the same monolayer to reduce data variability. Bradyzoites and tachyzoites were independently isolated (as above) and combined at a ratio of approximately 1:1. Each biological replicate challenged a panel of host cells with the same mixture of tachyzoites and bradyzoites; HFF cells were always included as a control. Cells were grown on 18 mm coverslips pre-coated with poly-l-lysine (0.01%) until confluent. For MDCK, EA.HY926, HCT-8 and CACO-2 cells, formation of tight junctions was confirmed by immunofluorescence assays against ZO-1 (see below).

Each cell type was challenged with 1ml parasites (0.4 – 1.1 × 10^5^ total parasites, 1-3 parasites per cell) for 1h at 37°C in 5% CO2. The exact starting ratio of bradyzoites to tachyzoites was then quantified by counting the number of RFP-(tachyzoites) and GFP-(bradyzoites) expressing parasites. After the 1 hr incubation, the monolayers were gently washed with 1ml PBS, fixed with 2% PFA for 15 minutes, and blocked for 30minutes with PBS containing 1% (w/v) bovine serum albumin (PBS-BSA). Extracellular parasites were stained by incubation for 60 min with anti-SAG3 mouse monoclonal antibody (#NR-50257, BEI Resources) diluted 1:500 in PBS-BSA. After washing 3 times with PBS-BSA, coverslips were incubated with either Alexafluor 405nm-or Alexafluor 647nm-conjugated anti-mouse secondary antibody (Invitrogen), diluted 1:500 in PBS-BSA, in the dark for 1 hour. To quantify the number of invaded parasites, 10 fields of view were imaged per biological replicate. Intracellular parasites were identified as Alexafluor 405/647 negative and positive for either RFP fluorescence (540nm excitation/605nm emission; tachyzoites) or GFP fluorescence (488nm excitation/534nm emission; bradyzoites). The relative number of invaded tachyzoites and bradyzoites parasites was then normalized using the measured ratio of tachyzoites and bradyzoites in the starting suspension and presented as normalized compared to invasion into HFFs.

### Transmigration

MDCK, HCT-8 or CACO-2 cells were grown to confluence on 3μm ThinCert™ Cell Culture Inserts (Griener bio) in 12 well plates. Confluence and monolayer polarization were confirmed prior to experiments by measuring transcellular electrical resistance (TCER) and performing immunofluorescence assays on the tight junction protein ZO-1 (data not shown); barrier integrity is reached after TCER plateaus and stabilizes at >2000 Ω/cm^2^ and >125 Ω/cm^2^ for MDCK/HCT-8 and CACO2 cells respectively ^[35-37]^. Stable TCER was confirmed for 2 days prior to challenge with parasites. In each biological replicate, cells were challenged with either no parasites (unchallenged), tachyzoites (0.5 – 1 × 10^6^), or bradyzoites (0.1 – 0.5 × 10^6^) for 6 hours. TCER measurements were made for each monolayer immediately after addition of parasites, and 6 hours later. In a separate experiment, where parasite transmigration was not quantified, tachyzoites (0.5 – 1 × 10^6^), or bradyzoites (0.1 – 0.5 × 10^6^) and FITC-dextran (MW 3000 Da, 1 mg/ml, Molecular Probes) in PBS were added to cell monolayers grown on ThinCert™ Cell Culture Inserts. The passage of FITC-dextran across the monolayer was quantified after 6 hours using a Biotek synergy 2 plate reader. The relative fluorescence intensity (RFI) of the dextran that had crossed the cell barrier was divided by the RFI of the dextran added to the top of the monolayer and multiplied by 100 to quantify the percentage of the added dextran that had crossed at some time within the 6-hour challenge. One monolayer for each cell type per experiment was treated with 2μg/ml CytoD to disrupt the tight junctions, resulting in reduced TCER and rapid flux of dextran and parasites across the cells and into the lower chamber of the well ^[38]^.

To quantify transmigration across the monolayers, the number of parasites/μl in the challenge suspension and in the lower chamber of the well 6-hours after challenge was quantified using a Miltenyl MACSQuant VYB. Bradyzoites and tachyzoites were distinguished based on RFP (PRU-RFP tachyzoites) or GFP (PRU-LDH2 GFP bradyzoites) fluorescence, respectively. The proportion of bradyzoites and tachyzoites able to transmigrate was normalized based on the numbers of parasites in the challenge suspension and expressed as a percentage of those added to the upper chamber.

### Egress

Infected cells containing mature tachyzoite vacuoles (∼48 hours post invasion) or bradyzoite cysts (5 – 6 days post differentiation) were incubated with 1μM Ionomycin or 1μM A23187 to stimulate egress. For tachyzoites, monolayers were fixed 2.5, 5- and 30-minutes after stimulation by gently replacing the media with 2% PFA in PBS for 10 minutes. For bradyzoites, monolayers were fixed 5-, 30-, 60- and 120-minutes following stimulation by gently replacing the media with 2% PFA in PBS for 10 minutes. For each biological replicate 100 vacuoles were scored as either intact or egressed based on the dispersal of tachyzoites (PRU RFP) or bradyzoites (PRU-LDH2 GFP).

## Results

### Type II (PRU) tachyzoites are as motile as type I (RH) tachyzoites

3D motility has only been characterized in Type I (RH strain) tachyzoites and mutant lines made in this background ^[20, 39-42].^ However, Type II strains (*e*.*g*., PRU) are much more efficient than Type I at generating bradyzoites, both *in vitro* and in mice, and therefore were preferred for a comparison of bradyzoite and tachyzoite motility. We first assessed how well PRU tachyzoites move in our established 3D motility assay ^[20]^. In contrast to what was previously observed in 2D motility assays with Type II Me49 tachyzoites ^[43]^, Type II PRU tachyzoites were as motile in the 3D motility assay as Type I RH tachyzoites. We saw no differences in the proportion of PRU *vs*. RH tachyzoites moving in 3D, the distance they moved (both first-to-last point displacement and track length), or their maximum and average speeds along the trajectory (Supplementary Figure 1). Thus, type II PRU strain parasites can serve as a good model to directly compare bradyzoite and tachyzoite motility.

### Bradyzoites are as motile as tachyzoites in a 3D model extracellular matrix

As bradyzoites and tachyzoites are morphologically indistinguishable in our 3D motility assay we utilized the PRU-derived fluorescent parasite reporter line, LDH2::GFP, in which GFP placed under the control of the *ldh2* stage-specific promoter is expressed exclusively in bradyzoites ^[44]^. Using this line allows us to specifically identify and track bradyzoites ^[20]^ by their GFP expression. Motility of the non-fluorescent tachyzoites is tracked by staining the nucleus with Hoechst 33342 (as in Supplementary Figure 1 and ref [20]).

First, we evaluated the motility of bradyzoites isolated from *in vitro* cultures where tachyzoite-to-bradyzoite differentiation was induced by standard stress conditions (pH 8.2, ambient CO_2_) ^[34]^. We compared the well characterized 3D motility of *in vitro-*derived tachyzoites to that of bradyzoites exposed to stress for five days. The bradyzoites were liberated from their surrounding cyst wall and host cell by mechanical disruption either with or without a one-minute pre-digestion with acid pepsin (+AP and -AP, respectively; Figure 1). Using the 3D motility assay to analyze hundreds of parasites, we found no significant difference in the ability of tachyzoites and bradyzoites isolated with AP to move (compare the two left panels and in Figure 1A and first and third bars in Figure 1B). Without AP digestion, we recovered only 30% the number bradyzoites recovered with AP, presumably due to retention within the cyst, and significantly fewer of the -AP bradyzoites moved (compare second and third bars in Figure 1B). We then compared the trajectory characteristics of those tachyzoites and bradyzoites (+AP, -AP) that moved. There were no significant differences between the tachyzoite and two bradyzoite preparations in: parasite displacement (Figure 1C, first three bars); track length (Figure 1D), and maximum (Figure 1E) and mean (Figure 1F) speeds achieved. Bradyzoites can therefore move as effectively as tachyzoites in 3D, and a brief digestion of the cyst wall by acid pepsin yields more motile bradyzoites from *in vitro* culture than mechanical disruption alone.

**Figure 1.**
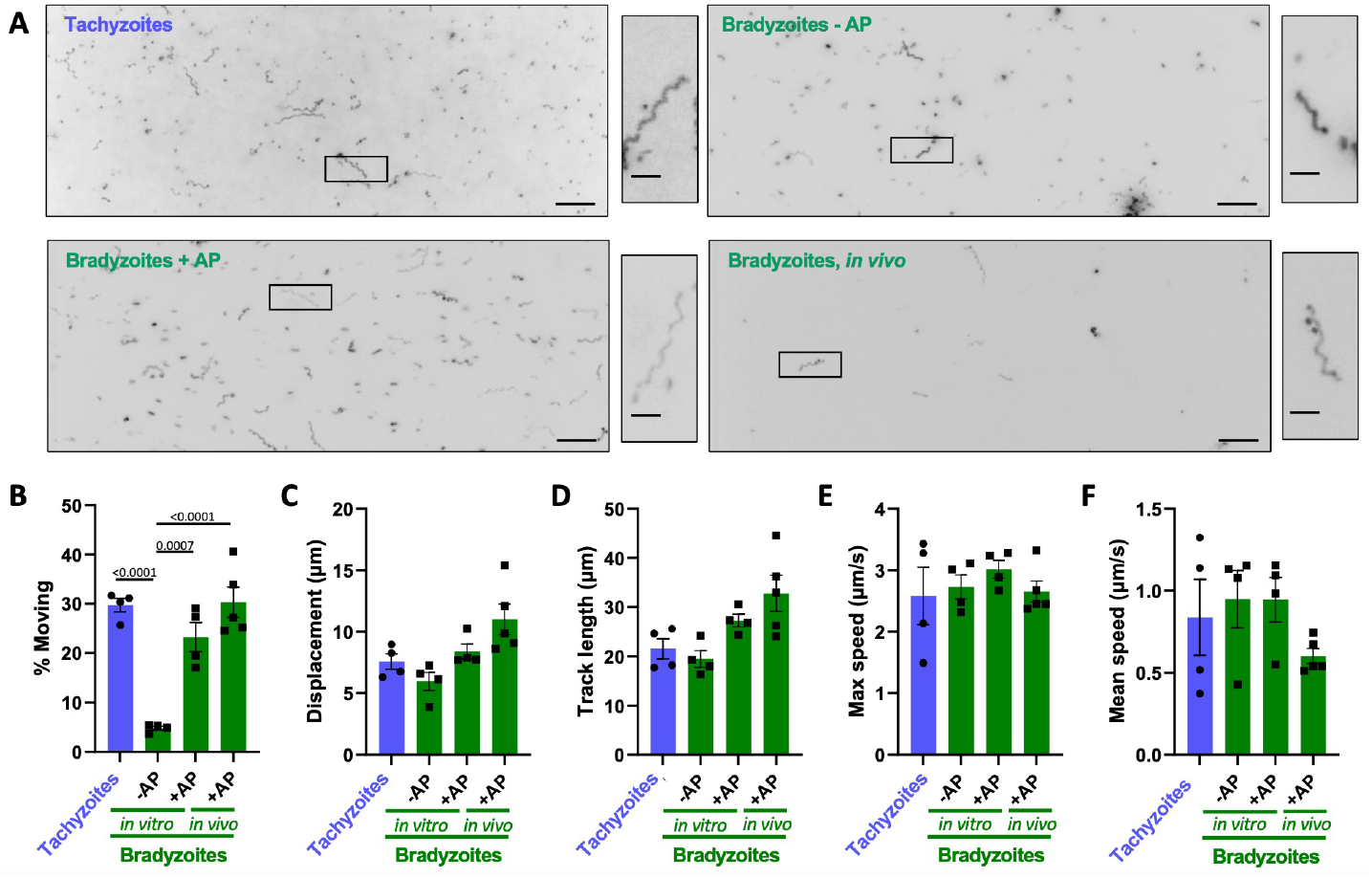
Comparison of the motility of PRU tachyzoites and bradyzoites. (A) Representative maximum intensity projections of parasites moving through Matrigel during 60 seconds of imaging; scale bar = 40μm. Insets (black boxes) are magnified, rotated and displayed to the right of each full field of view; scale bar 10μm. (B) Percentage of tachyzoites moving with displacements > 2.5μm and bradyzoites moving with displacements > 2.8μm during 60 seconds of imaging. (C-F) For all parasites that exceeded the 2.5/2.8 μm displacement thresholds (see Methods), the following median trajectory parameters were quantified: C) displacement; D) track length; E) maximum speed achieved across the entire track; and F) mean speed. On the graphs, each data point represents a biological replicate consisting of 3 – 6 technical replicates. Bar height shows the mean and error bars show the s.e.m. of the biological replicates. The number of parasites analyzed in B-F was 4387 (Tachyzoites), 1132 (Bradyzoites – AP), 3757 (Bradyzoites + AP), and 728 (Bradyzoites, *in vivo*) stage- and isolation method-specific differences in the motility parameters were compared using an ordinary one-way ANOVA with Tukeys correction for multiple comparisons; only statistically significant differences *(p*<0.05) are shown.

To ascertain if the motility of our *in vitro-*derived bradyzoites is representative of mature bradyzoites found in cysts *in vivo*, we also characterized the motility of bradyzoites isolated from the brains of chronically infected mice. The 3D motility assay requires parasite preparations relatively free of cellular contamination. Isolation of free bradyzoites from mouse brain to a sufficient level of purity is inefficient and yields are low: the number of bradyzoites isolated from the brains of 2 mice is equivalent to ∼4% of the number that can be obtained from one T75 culture flask of *in vitro-*differentiated parasites. Nevertheless, we recovered sufficient numbers of bradyzoites from the mouse brain to compare to *in vitro-*differentiated bradyzoites liberated from cysts (+AP). We see no significant difference in the proportion of parasites that move (Figure 1A, bottom two panels, and Figure 1B, right two bars). Comparing how far and fast these two bradyzoite populations move also reveals no significant differences in their median displacement, track length, or maximum and mean speeds (Figures 1C-F).

In addition to quantifying and comparing the average motility parameters, we analyzed the distributions of each of the motility parameters in the different parasite preparations to determine if the behavior of parasites across the population was consistent. We compared the 5% to 95% range that describes the motility of 90% of all parasites characterized and found no significant differences within this range for displacement, track length, or maximum and mean speeds (Supplementary Figures 2A-D).

These data demonstrate that, using the isolation method reported here, bradyzoites are motile and their motility is equivalent, in all aspects measured, to tachyzoites. Furthermore, 5-day stress-induced bradyzoites from *in vitro* culture (+AP) faithfully recapitulate the motility of bradyzoites isolated from the brains of infected mice, providing a more abundant source of parasites for studies of bradyzoite motility without having to use an animal model.

### Bradyzoites invade intestinal cells to a greater extent than tachyzoites

Because parasite motility plays a critical role in host cell invasion, we next determined if excysted bradyzoites were able to invade host cells as effectively as tachyzoites and if either stage showed host cell specificity. To this end, we developed a co-invasion assay in which approximately equal numbers of PRU-RFP tachyzoites ^[45]^ and PRU-LDH2::GFP bradyzoites ^[44]^ are mixed and used to challenge the same monolayer. Extracellular parasites are identified and excluded from quantification by staining the non-permeabilized samples for surface protein SAG3, which is expressed by both tachyzoites and bradyzoites ^[46]^. The cell types tested were: HFFs (human foreskin fibroblasts, the cells used to culture the parasites); EA.hy926 (human vascular endothelial) cells; MDCK (Madin-derby canine kidney) cells; and two intestinal epithelial cell lines, CACO-2 and HCT-8.

Compared to their levels of invasion into HFF and EA.hy926 cells, tachyzoites showed reduced invasion into CACO-2 and HCT-8 cells (50% and 60% respectively) and much reduced invasion into MDCK cells (∼7.5%) (Figure 2). With bradyzoites, invasion into MDCK cells is also reduced (∼8%) compared to HFF cells but, in contrast to tachyzoites, bradyzoites invade CACO-2 and HCT-8 cells at levels comparable to HFFs (Figure 2). Thus, bradyzoites invade intestinal epithelial cell monolayers more than twice as efficiently as tachyzoites, while no significant differences in invasion were observed for the other cell types tested.

**Figure 2.**
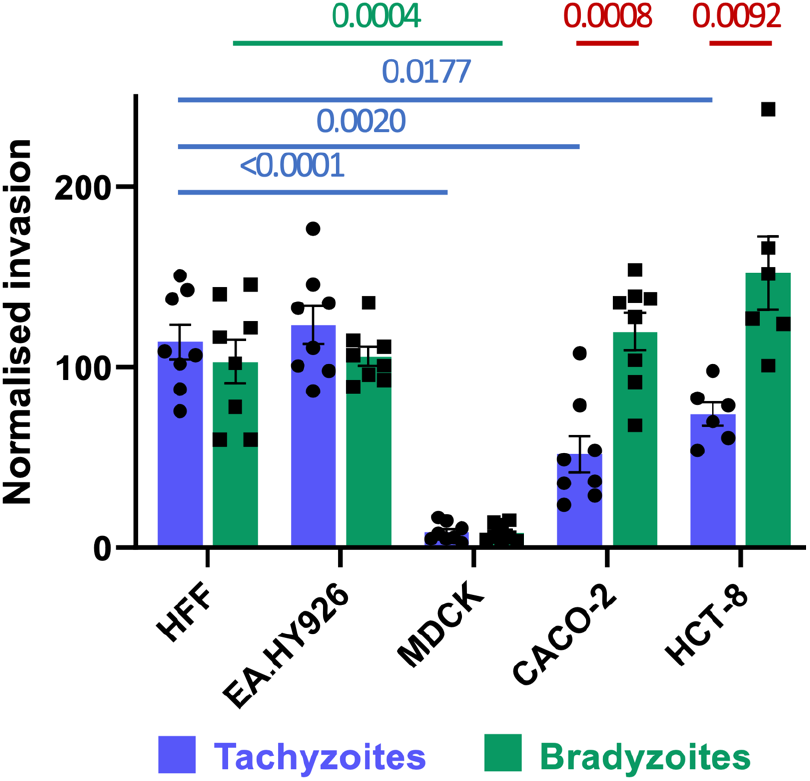
Impact of parasite stage and host cell type on invasion efficiency. Tachyzoite (blue) and bradyzoite (green) invasion into different cell types was determined by co-invasion assay. The data were normalized to the number of each parasite stage added to the well and the mean invasion into HFF cells. Invasion efficiency of tachyzoites (blue bars) and bradyzoites (green bars) into each cell type was compared to invasion into HFFs using an ordinary one-way ANOVA with Tukeys correction for multiple comparisons; significant differences are noted above the graph (blue lines, tachyzoites; green lines, bradyzoites). Invasion by the different stages into the same cell type was compared by unpaired students t-tests; significant differences are noted above the graph (red lines).

### Bradyzoites transmigrate across MDCK but not CACO-2 polarized monolayers more efficiently than tachyzoites

Establishing infection in the gut involves either invasion of and replication within intestinal epithelial cells or transmigration across the gut epithelium to access the lamina propria directly ^[43, 47]^. Having shown that tachyzoites and bradyzoites invade intestinal epithelial cells with differing efficiencies, we then tested whether there are differences in their ability to transmigrate across epithelial cell monolayers. First, we developed a transmigration assay that directly quantifies transmigration across the monolayer without coupling transmigration to subsequent host cell invasion as previously described ^[43, 47]^. We established a robust gating strategy to distinguish PRU-RFP tachyzoites and PRU-LDH2::GFP bradyzoites by flow cytometry (Figure 3A), and quantified the number of parasites (PRU-RFP tachyzoites, PRU-LDH2::GFP bradyzoites) that were able to transmigrate across an intact polarized monolayer of MDCK, HCT-8, or CACO-2 cells grown on ThinCert™ tissue culture inserts. The number of parasites recovered in the bottom well of the plate was then normalized to the number of parasites added to the monolayer, as quantified by flow cytometry, to compare transmigration efficiency.

**Figure 3.**
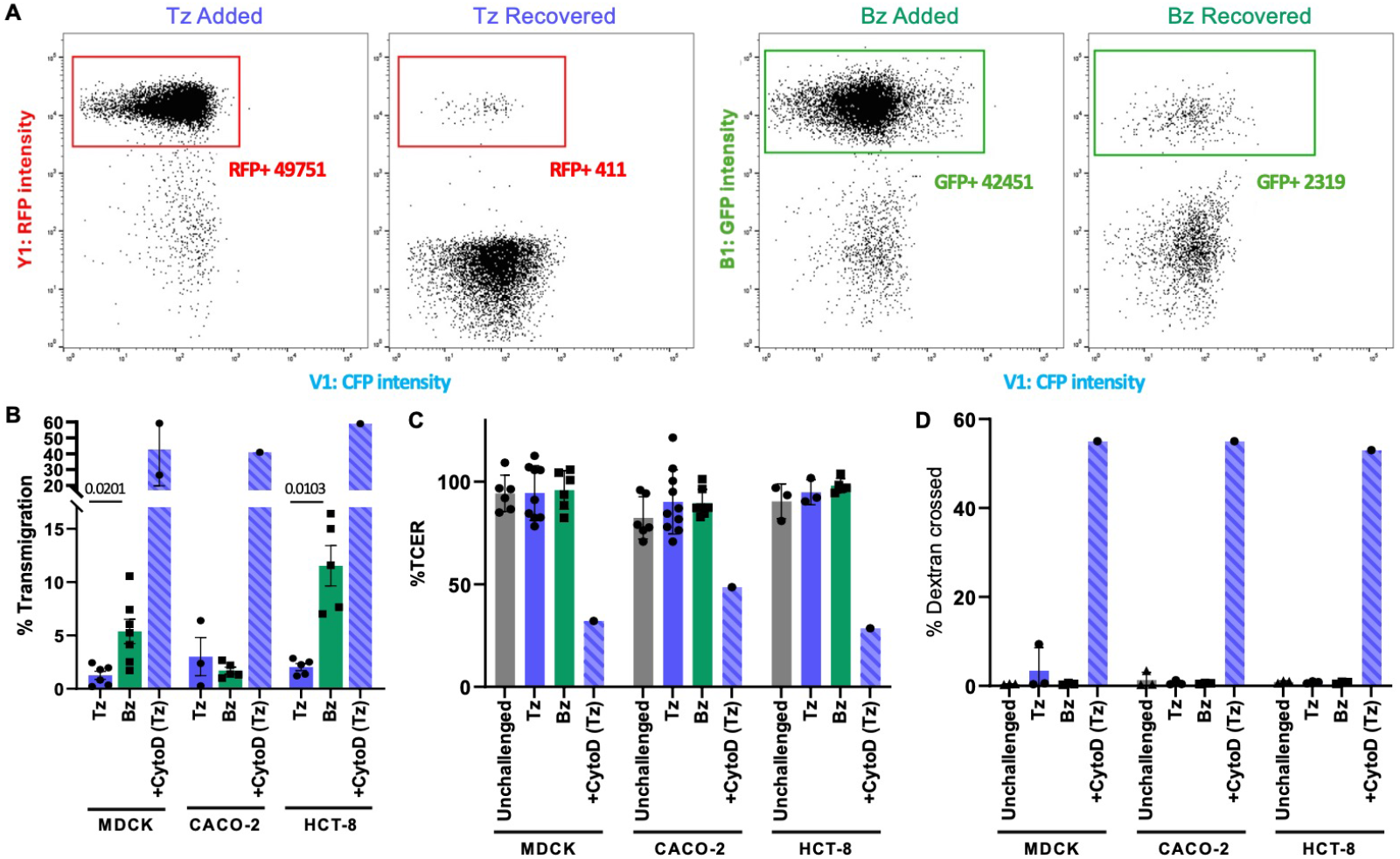
Bradyzoites transmigrate more efficiently than tachyzoites across MDCK and HCT-8 polarized monolayers, without altering barrier integrity. (A) Representative flow panels showing the gating strategy used to identify bradyzoites (left Bz GFP+) and tachyzoites (right Tz, RFP+). The strategy is used to quantify the number of parasites added to (Added) and transmigrated across (Recovered) a CACO-2 cell monolayer in 6 hours. (B) Quantification of the number of parasites that had crossed MDCK (left), CACO-2 (middle) or HCT-8 (right) monolayers within 6 hours of challenge. Addition of 2μg/ml cytochalasin D prior to tachyzoite challenge (CytoD Tz) disrupted the tight junctions of the monolayers and allowed the parasites to equilibrate between the upper and lower chambers. Tachyzoite vs bradyzoite transmigration was compared for each cell line using a students t-test; only significant differences are indicated. (C) The transcellular electrical resistance (TCER) prior to and 6 hours after challenge revealed no change due to parasite addition, unless the monolayers were intentionally disrupted with CytoD. The data were normalized to the average TCER measurement for all replicates. TCER values were compared using an ordinary one-way ANOVA and Tukeys correction for multiple comparisons. No significant differences were detected except in the samples that were treated with CytoD. (D) Cumulative dextran flux across the monolayers during the entire 6 hour challenge was quantified to determine if any transient changes in barrier integrity occurred. Dextran was only detected in the lower chamber when the monolayers were intentionally disrupted with cytoD. The data were analyzed using an ordinary one-way ANOVA and Tukeys correction for multiple comparisons.

Bradyzoites exhibited significantly greater transmigration across MDCK and HCT-8 monolayers than tachyzoites (4- and 5.5-fold increases in transmigration efficiency, respectively; Figure 3B). In contrast, we saw no significant difference between the number or bradyzoites and tachyzoites crossing a CACO-2 monolayer (Figure 3B). To determine if either stage compromised the barrier integrity of the monolayers, we monitored transepithelial electrical resistance (TCER) and saw no measurable decrease in resistance after 6 hours of challenge (Figure 4C) indicating no loss of barrier integrity when challenged with either bradyzoites or tachyzoites. As a positive control for barrier disruption, we pretreated monolayers with 2μg/ml cytochalasin D (CytoD) to disrupt tight junction integrity ^[38]^, prior to addition of the parasites. As expected, CytoD treatment led to rapid depolarization of the monolayer (Figure 3C) and increased passage of parasites across the monolayer (Figure 3B). We further confirmed that the barriers had not become transiently leaky during the 6-hour assay window by monitoring dextran flux across the monolayers. We saw no passage of fluorescent dextran from the top to the bottom well throughout the course of the experiment, while in monolayers treated with CytoD dextran fluorescence reached equilibrium between the two sides of the cell monolayer within 6 hours (Figure 3D). These data confirm and build upon previous reports demonstrating the ability of tachyzoites to transmigrate across MDCK ^[43]^, CACO-2 ^[47]^ and m-ICcl2 ^[47]^ cell monolayers, and show that enhanced bradyzoite transmigration across MDCK and HCT-8 monolayers reported here does not result from a change in monolayer integrity.

**Figure 4.**
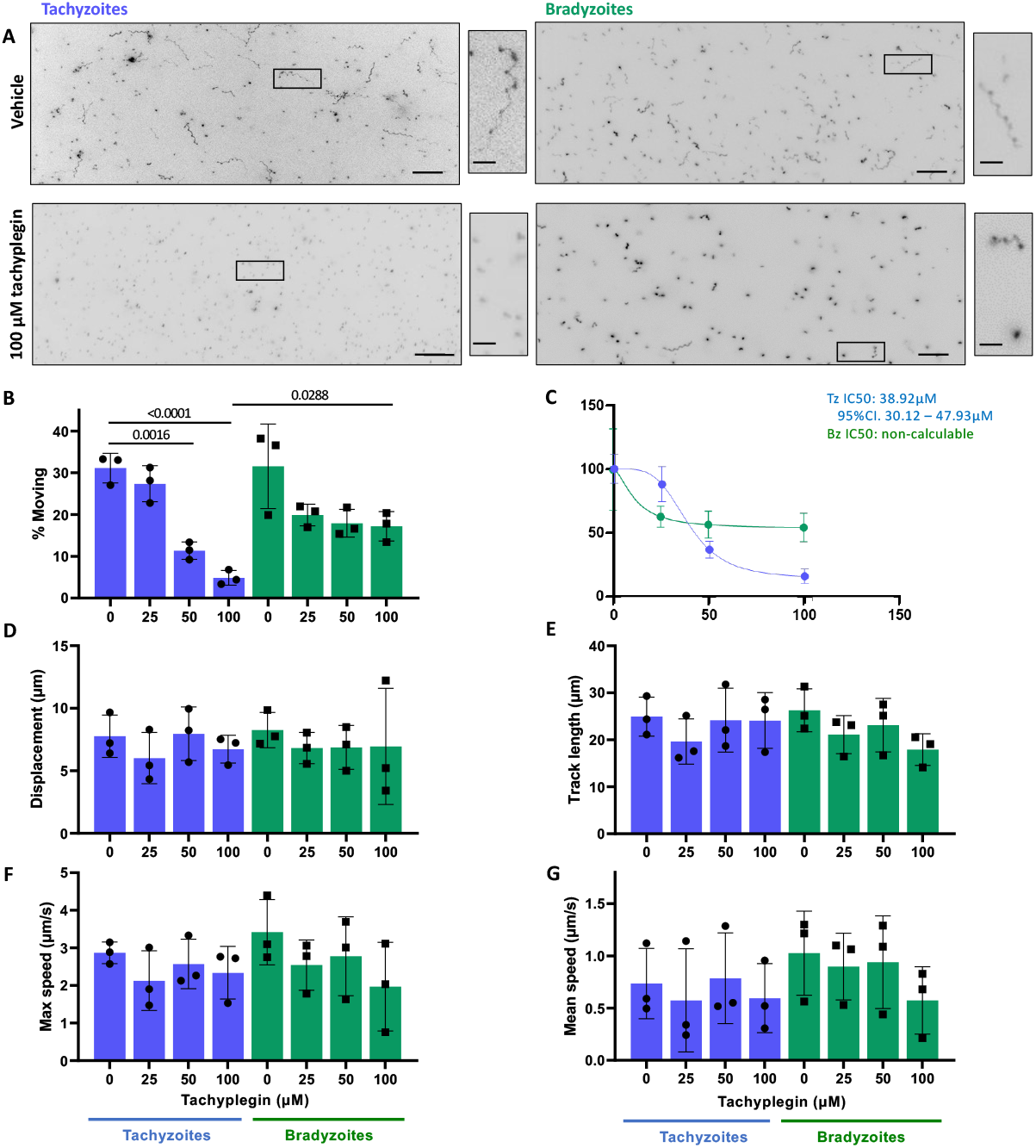
Comparison of tachyzoites and bradyzoite motility in the presence of tachyplegin. (A) Representative maximum intensity projections of tachyzoites and bradyzoites moving through Matrigel during 60 seconds of imaging in the absence (top 2 panels) or presence (bottom 2 panels) of 100μM tachyplegin; scale bar = 40μm. Insets (black boxes) are magnified, rotated and displayed to the right of each full field of view; scale bar 10μm. (B) Percentage of tachyzoites moving > 2.5μm and bradyzoites moving > 2.8μm during 60 seconds of imaging. (C) The IC50 for the % motility data shown in Panel A was calculated for tachyzoites (blue) to be 38.92μM; inhibition of bradyzoite motility (green) is insufficient to calculate an IC50. (D-G) For all parasites that exceeded the 2.5/2.8 μm displacement thresholds, the following median trajectory parameters were quantified: D) displacement; E) track length; F) maximum speed; and G) mean speed. On the graphs, each data point represents a biological replicate consisting of 2-3 technical replicates; top of the bars show the mean and error bars show the s.e.m. of the biological replicates. The number of parasites analyzed in B-G was 2043, 1552, 891, 755 (Tachyzoites 0, 25, 50, 100μM tachyplegin respectively) and 1265, 883, 1632, 1474 (Bradyzoites 0, 25, 50, 100μM tachyplegin respectively). The response of tachyzoites and bradyzoites at each concentration of tachyplegin was compared using unpaired students t-tests; a significant difference was observed only when motility of the two stages was compared for parasites treated with 100μM tachyplegin. The response of tachyzoites or bradyzoites to treatment with each concentration of compound was compared to the vehicle control (0) with an ordinary one-way ANOVA and Tukeys correction for multiple comparisons; only statistically significant differences (p<0.05) are shown.

### Pharmacological modulators of tachyzoite motility have both similar and distinctly different effects on bradyzoite motility

#### CytoD and KNX-002

We next wanted to determine if known modulators of tachyzoite motility have comparable effects on bradyzoite motility. CytoD inhibits actin polymerization, and has a well-documented inhibitory effect on tachyzoite motility in 2D ^[48]^. KNX-002 is a recently described inhibitor of the myosin motor TgMyoA that also inhibits tachyzoite motility ^[29]^. Given the central role of actin and TgMyoA in the motility of multiple apicomplexan zoites ^[24]^ we predicted that bradyzoite motility would also be inhibited by these compounds. This was indeed the case: in all measured parameters of motility and across all concentrations of compound, tachyzoites and bradyzoites were similarly inhibited by both CytoD and KNX-002 (Supplementary Figures 3 and 4, respectively). Quantification of the effect of the compounds on the proportion of parasites moving yielded consistent IC_50_ values of 0.12 and 0.10μM CytoD and 8.28 and 5.20μM KNX-002 for tachyzoites and bradyzoites, respectively, in each case with overlapping confidence intervals (Supplementary Figures 3C and 4C).

#### Tachyplegin

Tachyplegin is inhibitor of tachyzoite motility that targets myosin light chain-1 (MLC1), another critical component of the parasite’s myosin motor complex ^[21, 49, 50]^. Tachyplegin causes a steady dose-dependent decrease in the proportion of tachyzoites that move in 3D, over the range of 0-100μM compound (ref ^[49]^ and Figures 4A-C, tachyzoites). In contrast, while 25μM tachyplegin causes an initial decrease in the proportion of bradyzoites moving, increasing amounts of compound beyond 25μM do not inhibit motility further (Figures 4A-C, bradyzoites). As previously reported for RH strain tachyzoites ^[49]^, we saw no significant difference in any of the other calculated mean motility parameters in response to up to 100μM tachyplegin, in either tachyzoites or bradyzoites of the PRU strain (Figure 4D – G). For tachyzoites, treatment with tachyplegin also had no impact on the distributions of these motility parameters (Supplementary Figure 5). However, while bradyzoites treated with 100μM tachyplegin could still move, they were less able to move for long displacement distances than vehicle-treated parasites, as evident from the markedly truncated violin plot and a statistically significant difference in the displacement length for parasites in the 95^th^ percentile within each population (Supplementary Figure 5A). Thus, tachyplegin does not ablate bradyzoite motility to the same extent as tachyzoite motility at this higher dose but does reduce the distance the parasites can move (displacement) compared to untreated bradyzoites.

#### Enhancer 5

Enhancer 5 / Compound 130038 was previously shown to increase tachyzoite motility in 2D by stimulating intracellular calcium release, which leads to increased microneme secretion and TgMyoA phosphorylation ^[51, 52]^. 100μM enhancer 5 increased the proportion of tachyzoites moving in the 60 sec 3D motility assay, from 24% +/-3% to 52% +/-4%, but had no statistically significant effect on the proportion of bradyzoites moving (Supplementary Figure 6A and 6B). The effect of the enhancer is apparently specific to the proportion of tachyzoites moving: all other measured parameters of both tachyzoite and bradyzoite motility were unaffected by treatment with 100μM enhancer 5 (Supplementary Figure 6C-F).

#### Calcium ionophores

It has been previously reported that egress of Type II (Me49) tachyzoites from host cells is triggered by stimulation with calcium agonists, but egress of Me49 bradyzoites is not ^[27]^. We confirmed this observation with PRU parasites, showing that even extended (30 minutes) exposure to either ionomycin or A23187 caused no detectable egress of PRU bradyzoites compared to >80% egress by tachyzoites (Figure 5).

**Figure 5.**
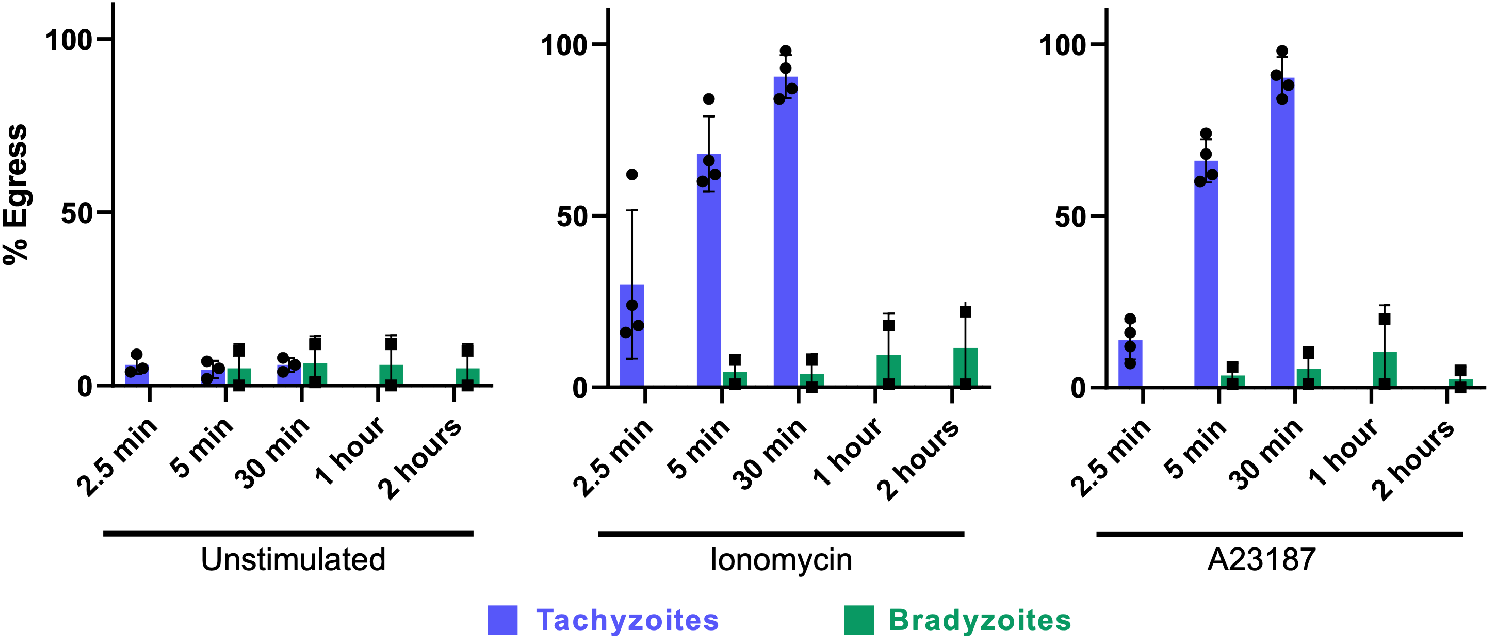
Bradyzoite egress is not calcium stimulated. Following treatment with the calcium agonists ionomycin (middle graph) and A23187 (right graph), tachyzoites (blue bars) rapidly leave the host cells in which they reside, whereas extended treatment with these agonists induces little to no bradyzoite (green bars) egress.

Taken together, these data demonstrate that while some pharmacological agents targeting the parasite’s motor machinery or the signaling mechanisms underlying motility have similar effects on tachyzoites and bradyzoites, other compounds have distinctly different effects on these two life-cycle stages.

## Discussion

Differentiation of *T. gondii* to bradyzoites and encystment in cells of the brain and other tissues results in a chronic infection. We show here that the bradyzoites within the cysts are primed to move upon excystation. Bradyzoites are as motile as tachyzoites when released from the cyst by the combination of mechanical shear and a short acid pepsin digestion and are at least as invasive as tachyzoites into a variety of cell types. The higher levels of invasion of bradyzoites relative to tachyzoites into CACO-2 and HCT-8 cells may reflect an enhanced ability of bradyzoites to invade cells of the gut epithelia. Bradyzoites also show an increased ability to transmigrate across polarized epithelial cell monolayers (MDCK and HCT-8 cells). Since establishment of an infection following ingestion of bradyzoite-containing cysts requires the parasites to rapidly leave the cyst, migrate to the gut wall and either actively invade into the gut epithelial cells or transmigrate between them to access the lamina propria, the enhanced ability of bradyzoites to invade gut cells and transmigrate across polarized epithelia may be adaptive traits in this parasite life cycle stage. Enhanced transmigration ability may be cell type-specific, however, as neither bradyzoites nor tachyzoites transmigrate efficiently across CACO-2 cells. Further investigation into stage-specific invasion into and transmigration across epithelial monolayers derived from complex mixtures of cell populations (*e*.*g*., open format enteroids ^[53]^) may reveal cell and gut regions that are particularly susceptible to bradyzoite infection, differences that homogeneous monolayers of cells fail to fully capture.

Two of the three tachyzoite motility inhibitors we tested (CytoD and KNX-002) show similar activity against bradyzoites and tachyzoites, suggesting conservation of an actin-myosin-based motility machinery between the two stages. Why then might tachyplegin, which targets a key component of this machinery – myosin light chain 1 (MLC1; ^[49]^) – have a different effect on tachyzoites and bradyzoites? One possibility is that the differential activity is not due to differences in the motor *per se*, but rather to differences in the phosphorylation of MLC1 in tachyzoites and bradyzoites. TgMLC1 is phosphorylated on at least nine distinct sites in tachyzoites ^[54]^; intriguingly, two of these phosphorylation sites (S55, S57) are located in close proximity to the tachyplegin binding site in the protein (C58) ^[49]^ and could conceivably affect compound binding. Furthermore, calcium-stimulated egress of tachyzoites is accompanied by the rapid phosphorylation of more than 50 proteins, including S55 of MLC1 ^[55]^. Perhaps the different signaling pathways that underlie tachyzoite egress and bradyzoite excystation ^[27]^ result in a different MLC1 phosphorylation profile and therefore a different sensitivity to tachyplegin. Further work will be required to test this specific hypothesis.

The work reported here confirms and extends previous work showing that the signaling underlying host cell egress by tachzyoites is different from the signaling underlying egress of bradyzoites from cysts. Not only are bradyzoites less responsive to calcium signaling within the cyst, but once released they are also less sensitive to the effects of enhancer 5. Enhancer 5 increases tachyzoite motility through an effect on calcium signaling and microneme secretion ^[52]^, but does not significantly increase bradyzoite motility in our 3D assay. This indicates that not only are encysted bradyzoites less responsive to calcium stimuli but recently excysted bradyzoites show dampened responses to calcium stimuli as well.

Taken together, the data presented here demonstrate that although the motility and motility-associated behaviors of tachyzoites and bradyzoites are similar in many respects, they also differ in ways that may be adaptive for that particular life cycle stage. Tachyzoites and bradyzoites also show differing sensitivity to compounds targeting the parasite’s motor machinery and the signaling processes underling its activation, and this needs to considered in drug development efforts ^[28-32]^ targeting *T. gondii* motility.

## Acknowledgements

We thank Bruno Martorelli di Genova, Frances Male, and Kevin Brown for helpful comments on the manuscript. Flow Cytometry was carried out in the Harry Hood Bassett Flow Cytometry Facility (RRID:SCR_022147) at the University of Vermont Larner College of Medicine; we thank the Director of the Facility, Dr. Roxana del Rio-Guerra, for assistance with the development of the flow-based transmigration assay. This work was supported by U.S. Public Health Service grant R01AI139201 (GEW) and American Heart Association grant 20POST35220017 (RK).

---

**Supplementary figure 1.**
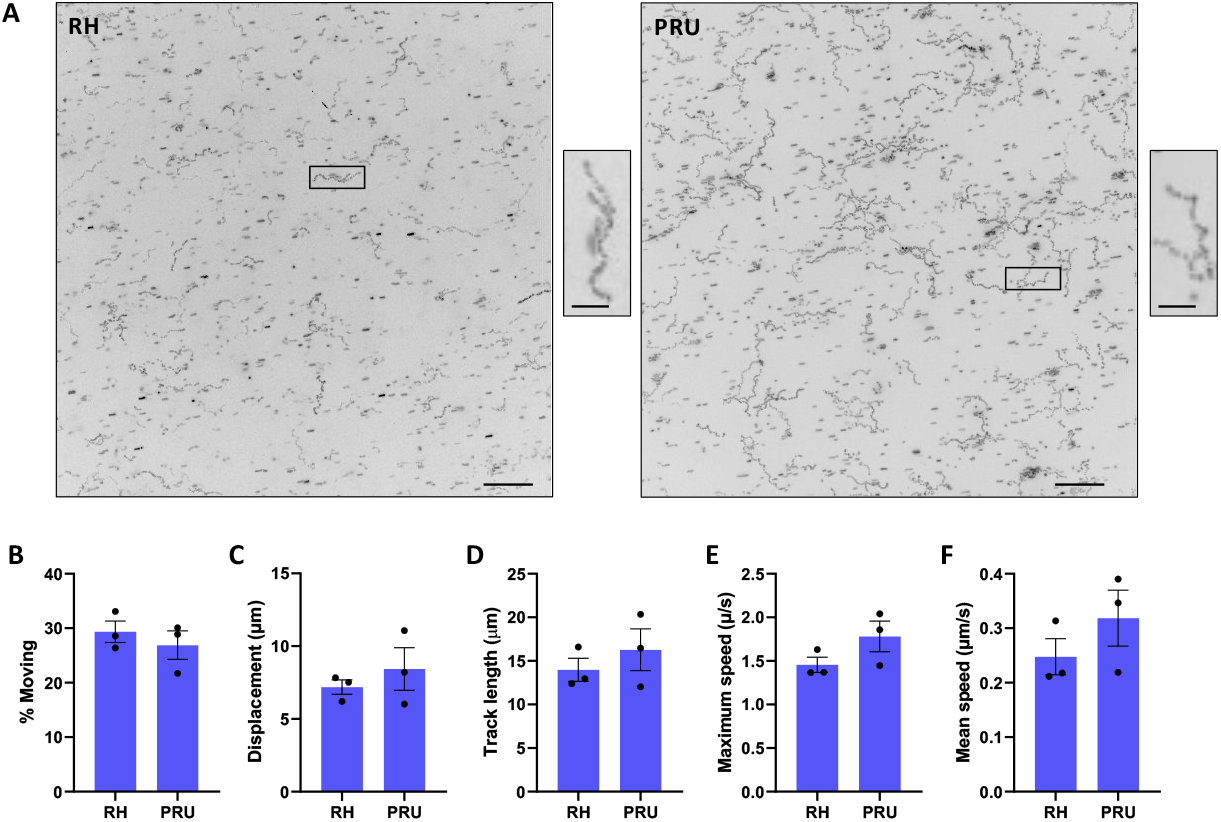
Comparison of RH and PRU tachyzoite motility. (A) Representative maximum intensity projections of RH (left panel) and PRU (right panel) tachyzoites moving through Matrigel during 60 seconds of imaging; scale bar = 40μm. Insets (black boxes) are magnified, rotated and displayed to the right of each full field of view; scale bar 10μm. (B) Percentage of parasites moving > 2μm during 60 seconds of imaging; total number of parasites analyzed = 9064 (RH) and 7788 (PRU). (C-F) For all moving parasites, the following median trajectory parameters were quantified: C) displacement (distance from first to last point); D) track length; E) maximum speed achieved across the entire track; and F) mean speed. Each pair of motility parameters (RH vs PRU) was compared using an unpaired students t-test, and no significant difference in these parameters was identified between the strains. On the graphs, each data point represents one of three biological replicates, each consisting of 2 – 4 technical replicates. Bar height shows the mean and error bars show the s.e.m. of the biological replicates.

**Supplementary figure 2.**
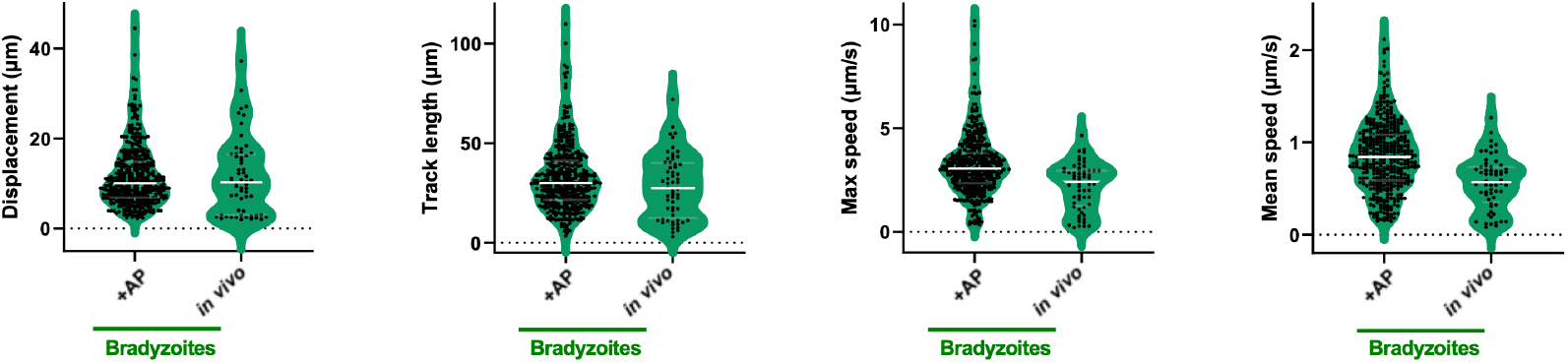
Comparison of the motility parameter distributions for *in vitro-vs. in vivo-*derived bradyzoites. (A) Displacement, (B) Track length, (C) Maximum speed and (D) Mean speed, for all motile parasites in the *in vitro* (plus acid pepsin digest) and *in vivo* populations, plotted as violin plots. The median of each distribution is shown as a horizontal white line. Using unpaired students t-tests; no significant differences were identified between the two populations for any of the four parameters when comparing the median, 5% or 95% values.

**Supplementary figure 3.**
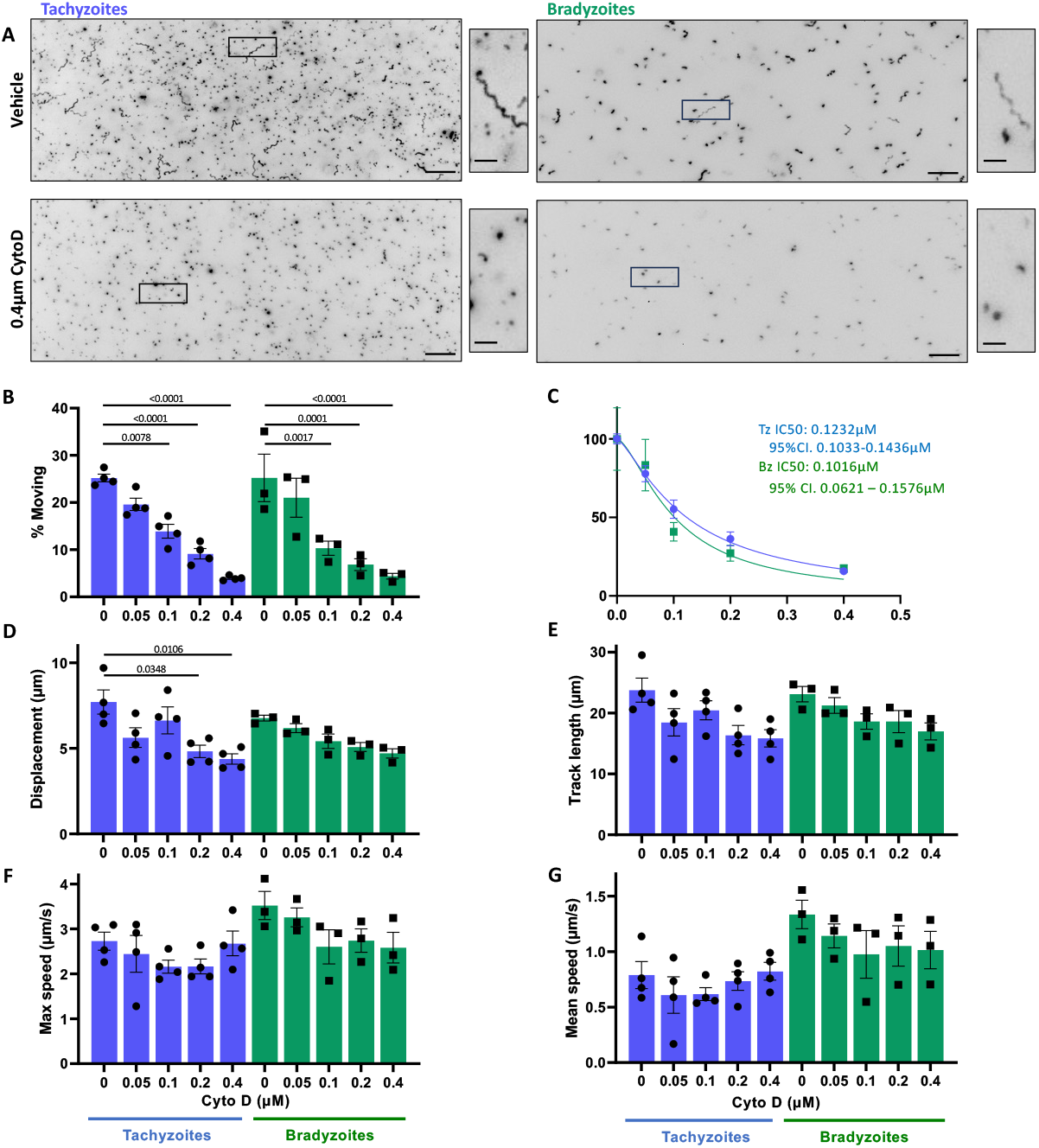
Comparison of tachyzoite and bradyzoite motility in the presence of Cytochalasin D. (A) Representative maximum intensity projections of tachyzoites and bradyzoites moving through Matrigel during 60 seconds of imaging in the absence (top 2 panels) or presence (bottom 2 panels) of 0.4μM Cytochalasin D (CytoD); scale bar = 40μm. Insets (black boxes) are magnified, rotated and displayed to the right of each full field of view; scale bar 10μm. (B) Percentage of tachyzoites moving > 2.5μm and bradyzoites moving > 2.8μm during 60 seconds of imaging. (C) The IC50 for the % motility data shown in Panel A was calculated for both tachyzoites (blue) and bradyzoites (green); no significant difference was seen in their response to treatment, as indicated by the overlapping 95% confidence intervals (CI). (D-G) For all parasites that exceeded the 2.5/2.8μm displacement thresholds, the following median trajectory parameters were quantified: D) displacement; E) track length; F) maximum speed; and G) mean speed. On the graphs, each data point represents a biological replicate consisting of 2-3 technical replicates; bar height shows the mean and error bars show the s.e.m. of the biological replicates. The response of tachyzoites and bradyzoites at each concentration of CytoD was compared using unpaired students t-tests; no significant differences were identified between the two stages at any dose. The number of parasites analyzed in B-G was 4129, 2440, 1086, 906 (Tachyzoites 0, 0.05, 0.1, 0.2, 0.4μM CytoD respectively) and 2529, 1220, 1786, 1109, 568 (Bradyzoites 0, 0.05, 0.1, 0.2, 0.4μM CytoD respectively). The response of tachyzoites or bradyzoites to treatment with each concentration of compound was compared to the vehicle control (0) using an ordinary one-way ANOVA and Tukeys correction for multiple comparisons; only statistically significant differences (p<0.05) are shown.

**Supplementary figure 4.**
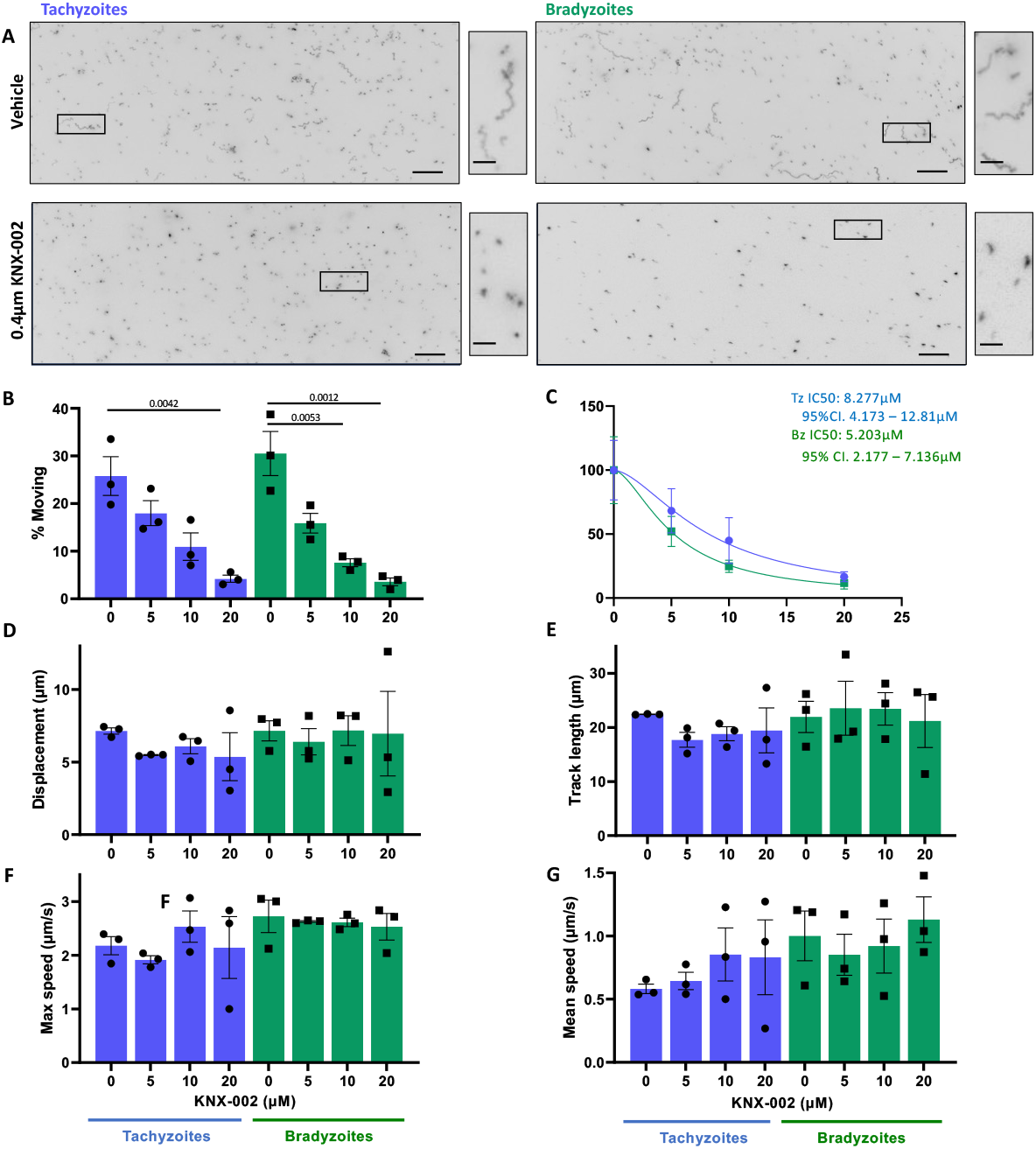
Comparison of tachyzoite and bradyzoite motility in the presence of KNX-002. (A) Representative maximum intensity projections of tachyzoites and bradyzoites moving through Matrigel during 60 seconds of imaging in the absence (top 2 panels) or, presence (bottom 2 panels) of 20μM KNX-002; scale bar = 40μm. Insets (black boxes) are magnified, rotated and displayed to the right of each full field of view, scale bar 10μm. (B) Percentage of tachyzoites moving > 2.5μm and bradyzoites moving > 2.8μm during 60 seconds of imaging. (C) The IC50 for the % motility data shown in Panel A was calculated for both tachyzoites (blue) and bradyzoites (green); no significant difference was seen in their response to treatment, as indicated by the overlapping 95% confidence intervals (CI). (D-G) For all parasites that exceeded the 2.5/2.8 μm displacement thresholds, the following median trajectory parameters were quantified: D) displac8ement; E) track length; F) maximum speed; and G) mean speed. For each graph, each data point represents a biological replicate consisting of 2-3 technical replicates; bar height shows the mean and error bars show the s.e.m. of the biological replicates. The response of tachyzoites and bradyzoites at each concentration of KNX-002 was compared using unpaired students t-tests; no significant differences were identified between the two stages at any dose. The number of parasites analyzed in B-G was 2112, 1186, 1251, 715 (Tachyzoites 0, 5, 10, 20μM KNX-002 respectively), and 3108, 1199, 868, 600 (Bradyzoites 0, 5, 10, 20μM KNX-002 respectively). The response of tachyzoites or bradyzoites to treatment with each concentration of compound was compared to the vehicle control (0) with an ordinary one-way ANOVA and Tukeys correction for multiple comparisons; only statistically significant differences (*p*<0.05) are shown.

**Supplementary figure 5.**
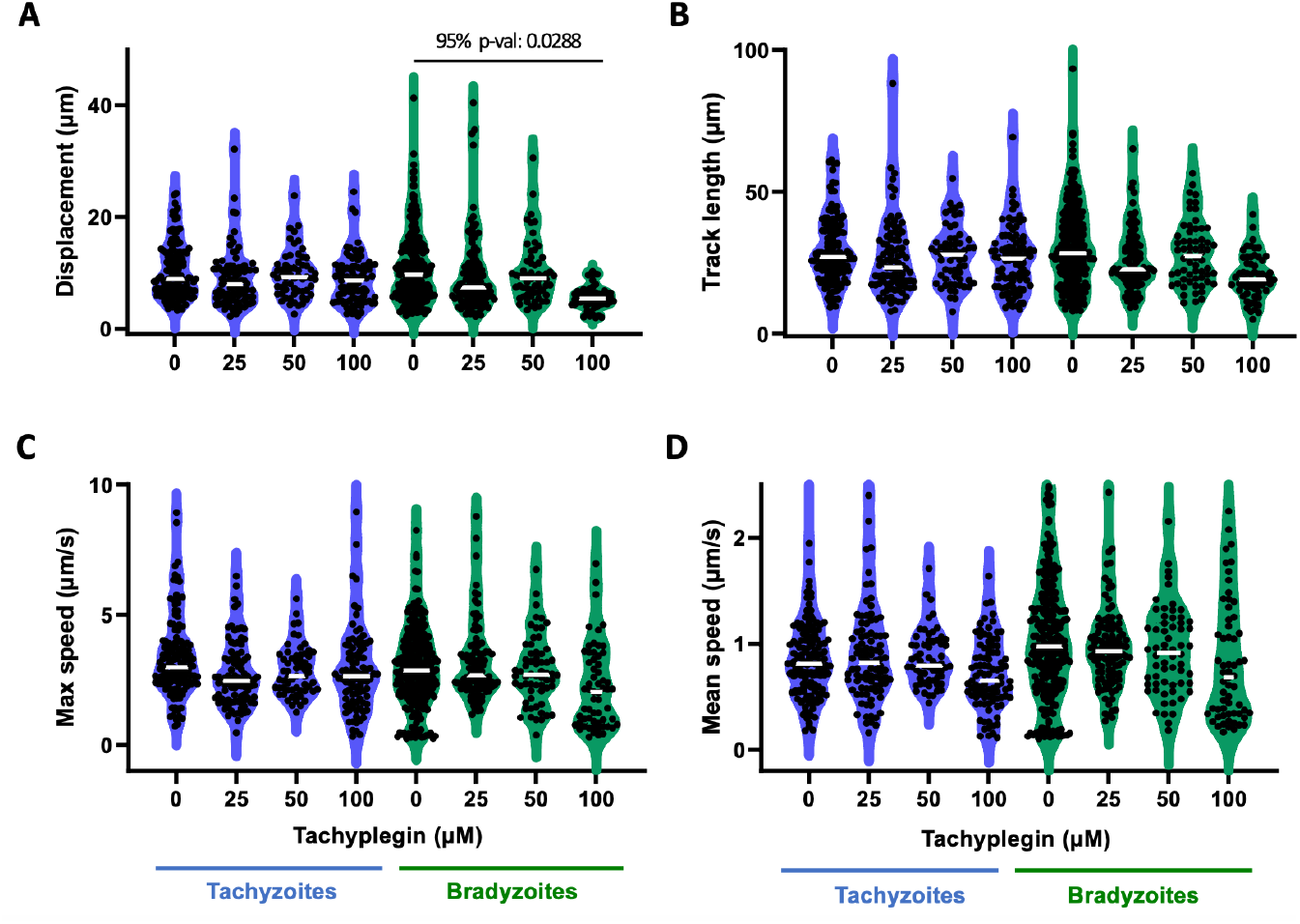
Comparison of the motility parameter distributions for tachyzoites and bradyzoites treated with tachyplegin. Tachyzoites and bradyzoites were treated with increasing doses of tachyplegin. The motility parameters compared were (A) displacement, (B) track length (C) maximum speed achieved and (D) mean speed. The 5^th^ and 95^th^ percentile values for each stage and compound concentration were compared to the vehicle control (0) with an ordinary one-way ANOVA and Tukeys correction for multiple comparisons. The only statistically significant difference was a decrease in the 95^th^ percentile of bradyzoite displacement, comparing 100μM tachyplegin to vehicle control.

**Supplementary figure 6.**
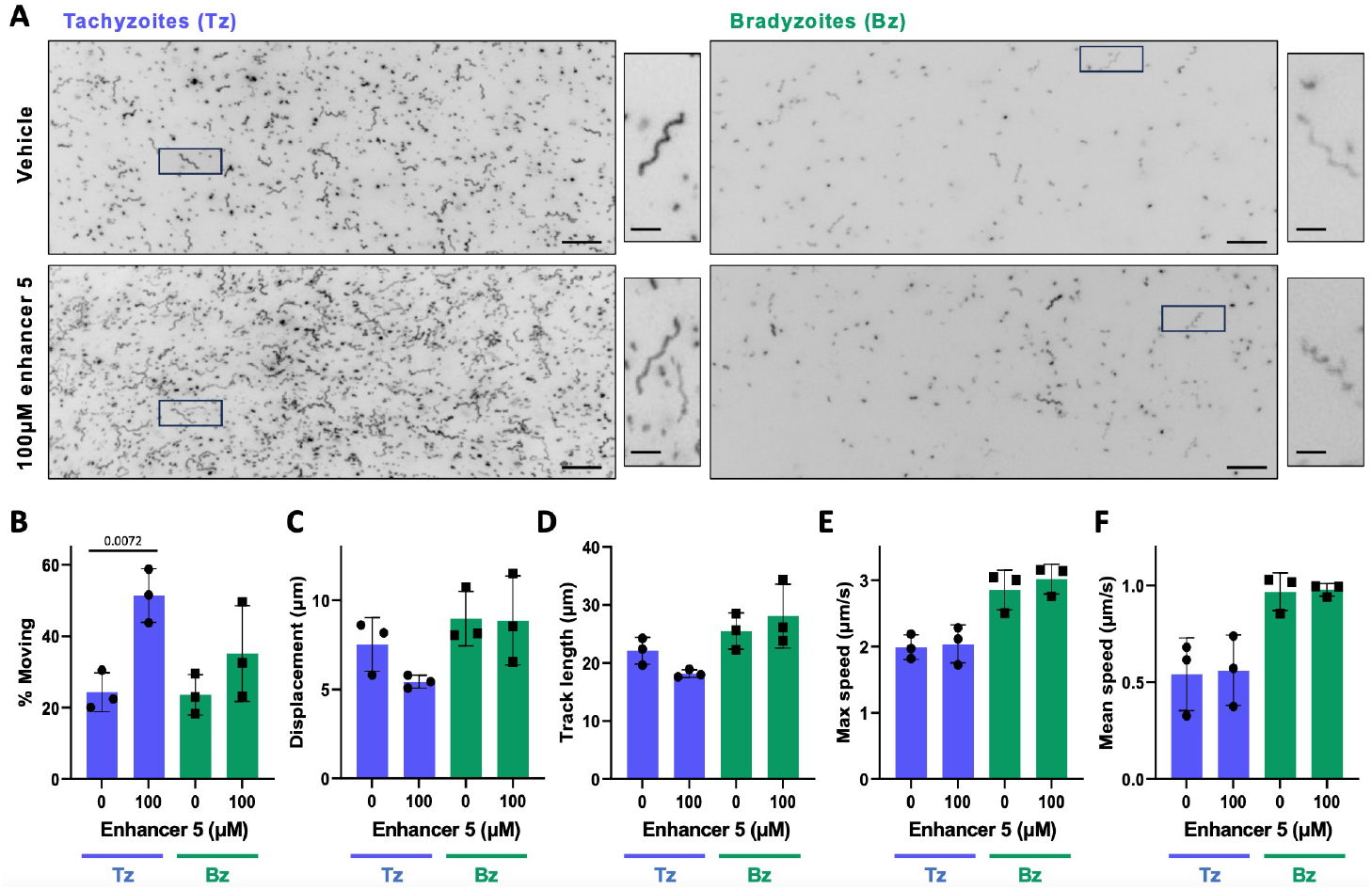
Comparison of tachyzoite and bradyzoite motility in the presence of enhancer 5. (A) Representative maximum intensity projections of tachyzoites (Tz) and bradyzoites (Bz) moving through Matrigel during 60 seconds of imaging in the presence of 100μMol enhancer 5; scale bar = 40μm. Insets (black boxes) are magnified, rotated and displayed to the right of each full field of view; scale bar 10μm. (B) Percentage of tachyzoites moving > 2.5μm and bradyzoites moving > 2.8μm during 60 seconds of imaging. (C-F) For all parasites that exceeded the 2.5/2.8 μm displacement thresholds, the following median trajectory parameters were quantified: C) displacement; D) track length; E) maximum speed; and F) the mean speed. Each data point represents a biological replicate consisting of 2-4 technical replicates; top of the bars show the mean and error bars show the s.e.m. of the biological replicates. The number of parasites analyzed in B-F was 2359, 2810 (Tachyzoites 0, 100μM enhancer 5 respectively) and 1743, 2109 (Bradyzoites 0,100μM enhancer 5 respectively). The response of tachyzoites and bradyzoites to enhancer 5 was compared to vehicle only (0) using unpaired students t-tests; only significant differences (*p*<0.05) are shown.

## Notes

### Competing Interest Statement

The authors have declared no competing interest.

